# Natural alleles of *LEAFY* and *WAPO1* interact to regulate spikelet number per spike in wheat

**DOI:** 10.1101/2024.08.17.608376

**Authors:** Junli Zhang, German F. Burguener, Francine Paraiso, Jorge Dubcovsky

## Abstract

Spikelet number per spike (SNS) is an important yield component in wheat that determines the maximum number of grains that can be formed in a wheat spike. In wheat, loss-of-function mutations in LEAFY (LFY) or its interacting protein WHEAT ORTHOLOG OF APO1 (WAPO1) significantly reduce SNS by reducing the rate of formation of spikelet meristems. In previous studies, we identified a natural amino acid change in WAPO1 (C47F) that significantly increases SNS in hexaploid wheat. In this study, we searched for natural variants in LFY that were associated with differences in SNS, and detected significant effects in the *LFY-B* region in a nested association mapping population. We generated a large mapping population and confirmed that the LFY-B polymorphism R80S is linked with the differences in SNS, suggesting that *LFY-B* is the likely causal gene. A haplotype analysis revealed two amino acid changes P34L and R80S, which were both enriched during wheat domestication and breeding suggesting positive selection. We also explored the interactions between the LFY and WAPO1 natural variants using biparental populations and identified significant interaction, in which the positive effect of the 80S and 34L alleles from LFY-B was only detected in the WAPO-A1 47F background but not in the 47C background. Based on these results we propose that the allele combination WAPO-A1-47F / LFY-B 34L 80S can be used in wheat breeding programs to maximize SNS and increase grain yield potential in wheat.

**Key message:** Specific combinations of *LFY* and *WAPO1* natural alleles maximize spikelet number per spike in wheat.

## Introduction

Wheat is an important global crop that contributes about 20 % of the calories and 24 % of the proteins consumed by the human population (D’Odorico et al. 2014). Therefore, continuous increases in grain yield are required to feed a growing human population. Grain yield is determined by the number of spikes per surface unit, the number of spikelets per spike, the number of grains per spikelet and grain weight. Among these yield components, spikelet number per spike (SNS) has the highest heritability because it is determined early during reproductive development and is not affected by later environmental conditions (Zhang et al. 2018).

The high heritability of SNS has facilitated its genetic dissection and several genes have been identified during the last five years affecting this trait. MADS-box transcription factors of the SQUAMOSA (*VRN1* and *FUL2*) and SVP clades (*SVP1* and *VRT2*) act as negative regulators of SNS, with loss-of-function mutations in any of these genes resulting in highly significant increases in SNS (*vrn1* 58%, *ful2* 10%, *vrt2* 20% and *svp1* 12%) (Li et al. 2019; Li et al. 2021). The combined *vrn1 ful2* mutants are unable to form spikelets and result in an indeterminate inflorescence meristem indicating that they are essential for the formation of a terminal spikelet (Li et al. 2019).

Several genes that modify heading time by regulating the expression of *VRN1* (Alvarez et al. 2023; Li et al. 2008; Li et al. 2015; Lv et al. 2014; Shaw et al. 2019; Turner et al. 2005; Yan et al. 2006) have been identified as negative regulators of SNS. Loss-of-function mutations in flowering promoting genes *PPD1* (Shaw et al. 2013), *FT1* (Chen et al. 2022) and *FT2* (Shaw et al. 2019) result in later heading and significant increases in SNS, whereas mutations in the *ELF3* flowering repressor (a negative regulator of *PPD1*) result in early heading and reduced SNS (Alvarez et al. 2023). The bZIPC1 protein physically interacts with FT2, and combined loss-of-function mutations in *bZIPC-A1* and *bZIPC-B1* in tetraploid wheat result in a large reduction in SNS with a limited effect on heading date (Glenn et al. 2023), but their effects on *VRN1* have not been reported.

A separate group of genes affect SNS mainly by repressing the formation of multiple spikelets per node (supernumerary spikelets). Mutations in *FZP* (Dobrovolskaya et al. 2015; Du et al. 2021), *TB1* (Dixon et al. 2018), or *DUO* (a positive regulator of the previous two genes) (Wang et al. 2022) result in the formation of supernumerary spikelets resulting in increased SNS.

Finally, three other genes have been identified that affect SNS without producing supernumerary spikelets. A rare allele of *COL5* from emmer wheat or its overexpression in transgenic plants was associated with increases in SNS, suggesting that this gene operates as a positive regulator of SNS (Zhang et al. 2022). This seems to be also the case for *WAPO1* and *LFY*, since null mutations in the two paralogs of these genes result in drastic reductions in SNS (Kuzay et al. 2022; Paraiso et al. 2024). Interestingly, the combined mutant with no functional copies of both genes showed similar SNS as the single mutants indicating a reciprocal recessive epistatic interaction between these two genes, and suggesting that *LFY* and *WAPO1* need each other to regulate SNS. This conclusion was also supported by the physical interaction between the two proteins in wheat protoplasts (Paraiso et al. 2024).

*WAPO-A1* was identified as the causal gene for a strong QTL for SNS in wheat (Kuzay et al. 2022; Kuzay et al. 2019), which was detected in multiple genome wide association studies (GWAS) (Muqaddasi et al. 2019; Voss-Fels et al. 2019). A natural allele resulting in a C47F amino acid polymorphism within the conserved F-box domain was associated with significant increases in SNS. The frequency of the 47F allele rapidly increased from tetraploid to hexaploid wheat and from old hexaploid landraces to modern hexaploid cultivars, suggesting positive selection by wheat breeders of this allele (Kuzay et al. 2019).

WAPO1 and its interacting partner LFY control SNS in wheat by affecting the rate of formation of spikelet meristems from the initial stages of spike development, rather than by controlling the timing of the transition between the inflorescence meristem and the terminal spikelet (Paraiso et al. 2024). The orthologous genes in rice (*ABERRANT PANICLE ORGANIZATION 1* (*APO1*) and *APO2*) also control the number of spikelets per panicle, show reciprocal recessive epistatic interactions for spikelet number, and encode proteins that physically interact with each other, suggesting a conserved role in the grasses (Ikeda-Kawakatsu et al. 2012; Ikeda et al. 2007).

In Arabidopsis, LFY also interacts with UNUSUAL FLORAL ORGANS (UFO, the ortholog of WAPO1) redirecting LFY to a different set of target genes (Rieu et al. 2023). LFY functions as a pioneer transcription factor that can access closed chromatin, playing an early role in different regulatory pathways (Jin et al. 2021; Lai et al. 2021; Yamaguchi 2021). Loss-of-function mutations in *LFY* or *WAPO1/UFO* result in the misregulation of floral homeotic genes and floral defects in the inner floral whorls in both Arabidopsis (Huala et al. 1992; Weigel et al. 1992) and in grass species (Bomblies et al. 2003; Ikeda-Kawakatsu et al. 2012; Paraiso et al. 2024; Selva et al. 2021), suggesting a conserved function in floral development. However, the role of LFY on floral meristem identity and regulation of inflorescence development is different in the grasses and Arabidopsis (Ikeda-Kawakatsu et al. 2012; Paraiso et al. 2024).

The floral defects associated with the loss-of-function mutations or the overexpression of both *WAPO1* and *LFY* result in reduced fertility, so they cannot be used in breeding. Instead, natural alleles go through selection, which eliminates mutations with negative pleiotropic effects on fertility or grain production. The identified natural 47F allele of *WAPO-A1* showed significant increases in SNS without affecting flower development or fertility. Based on the encouraging results from *WAPO-A1* and the known interaction between WAPO1 and LFY, we explored the natural variation in *LFY* to identify useful polymorphisms for wheat breeding. We identified two major amino acid polymorphisms associated with differences in SNS, and quantified their frequencies in modern and ancestral wheat germplasm. Finally, we investigated the epistatic interactions between the *LFY* and *WAPO1* natural alleles and identified combinations that maximizes SNS, which can be valuable for wheat breeding programs.

## Materials and Methods

### Plant Materials

We first explored the presence of QTL for SNS linked to the three *LFY* homeologs in hexaploid wheat in eight nested association mapping populations that were previously characterized for SNS in three field experiments under full and limited irrigation at the University of California, Davis between 2014 and 2016 (Zhang et al. 2018). We did not find significant associations between SNS and SNP markers within 10 Mb intervals at each side of *LFY-A* (*TraesCS2A02G443100*) or *LFY-D* (*TraesCS2D02G442200*), but we detected significant differences associated with the *LFY-B* (*TraesCS2B02G464200*) region in ‘Berkut’ x ‘PBW343’ population. To validate this association, we developed heterogeneous inbred families (HIFs) from two F_5_ lines (HIF40 and HIF63) from this population that were still heterozygous for the *LFY-B* region, and evaluated them for SNS.

To confirm the linkage between *LFY-B* and the QTL for SNS, we screened 402 F_5:3_ lines (804 gametes) for recombination events between markers 2B-657.1 (2B: 657,072,439) and 2B-660.1 (660,114,463, Table S1) flanking the *LFY-B* gene (TraesCS2B02G464200, chr2B: 658,198,819 to 658,201,834). We genotyped the recombinant lines with six additional markers developed within this region (Table S1) and identified three F_5:3_ lines heterozygous for *LFY-B* with close recombination events at both sides of *LFY-B* (Fig. 1C). From the progeny of these lines, we selected 10 F_7:2_ plants homozygous for the recombinant chromosome and 10 sister plants homozygous for the non-recombinant chromosome, carrying the alternative *LFY-B* alleles. We evaluated the F_7:3_ progeny of these 20 lines as single 1-m rows in the field in 2021, at the UC Davis Experimental Station.

**Fig. 1.**
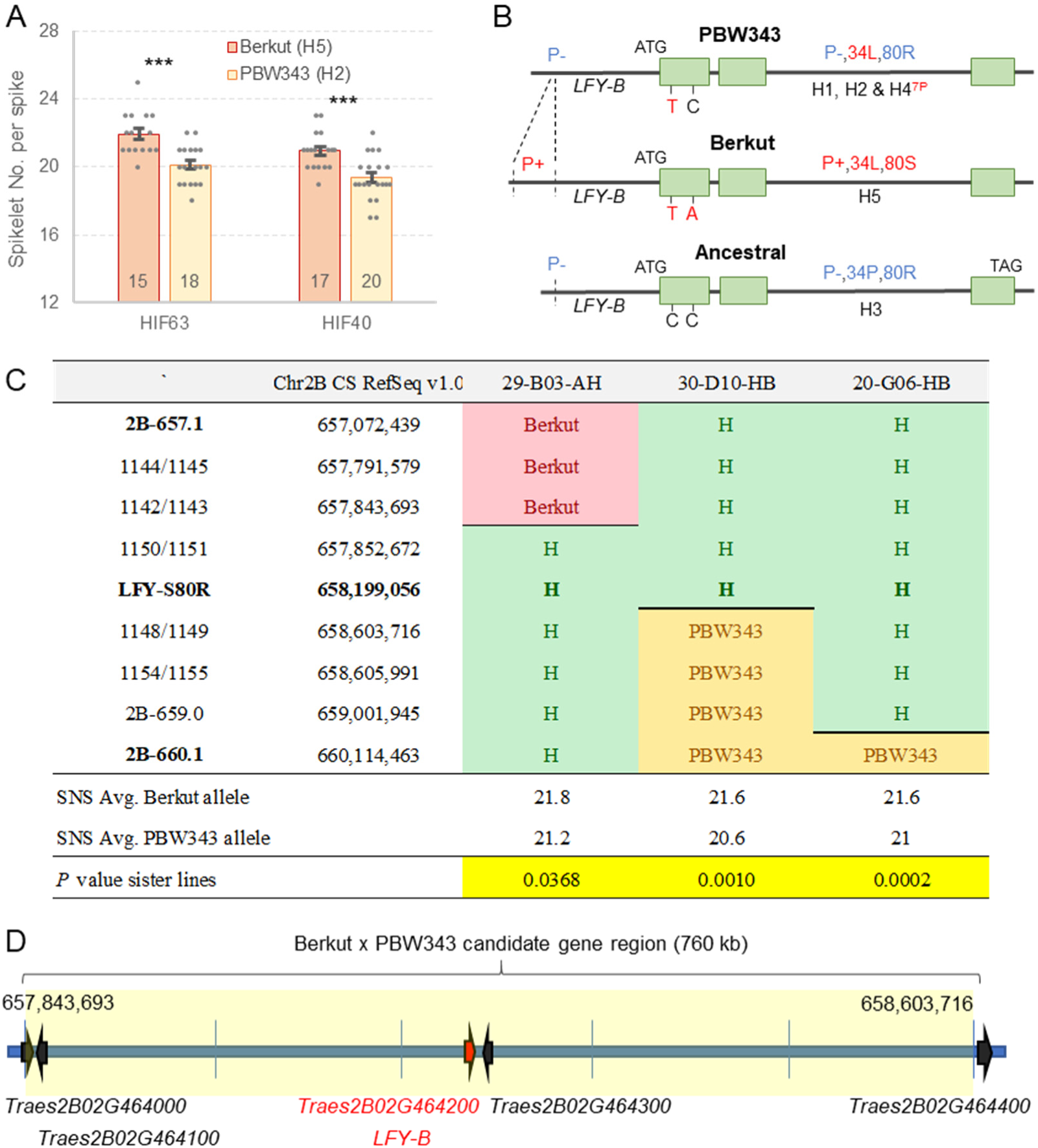
Polymorphisms in *LFY-B* are associated with changes in SNS in wheat. (**A**) Differences in SNS between sister HIF lines homozygous for the *LFY-B* alleles from PBW343 and Berkut in F_5:2_ heterogeneous inbred families (HIFs). Raw data and statistics are in Table S3. (**B**) *LFY-B* alleles differ in a promoter insertion (P+), amino acid changes P34L and R80S, and a proline deletion (8P to 7P, only in H4). Derived alleles are indicated in red. (**C**) Molecular markers and lines with the closest recombination events to *LFY-B.* SNS averages presented below correspond to progeny tests including lines homozygous for the recombinant and non-recombinant chromosome for each of the three families (n=10 1m-rows for each genotype in each of the three families). Significant differences indicate that the gene responsible for the differences in SNS maps to the central region linked with the *LFY-B* polymorphism. (**D**) Schematic representation of the genes detected within the 760 kb candidate gene region for the SNS QTL on chromosome arm 2BL.

### Natural variation and haplotype analysis

We performed a haplotype analysis for an 813.4 kb region of chromosome 2B including *LFY-B* (657,790,878 to 658,604,020 in CS RefSeq v1.1) using exome capture data from 56 common and pasta wheat accessions (https://wheat.triticeaetoolbox.org/downloads/download-vcf.pl) generated by the WheatCAP project (Exome_Capture_2017_WheatCAP_UCD). To obtain the complete *LFY* sequences associated with the different haplotypes, we assigned the different wheat sequenced genomes (Avni et al. 2017; Maccaferri et al. 2019; Walkowiak et al. 2020; Sato et al. 2021) to the haplotypes identified in the previous analysis.

The previous analysis revealed four polymorphisms among the different *LFY* alleles, for which we developed molecular markers (Table S1). We used these markers to screen a wheat germplasm collection including 85 *T. turgidum* ssp. *dicoccoides*, 86 *T. turgidum* ssp. *dicoccon*, 384 *T. turgidum* ssp. *durum*, and 234 *T. aestivum* accessions, which are part of a photoperiod insensitive spring wheat panel from North America previously evaluated for multiple agronomic traits (Zhang et al. 2018). We also checked the sequencing data of 35 accessions of Sitopsis species (*Ae. speltoides* (9), *Ae. sharonensis* (7), *Ae. longissima* (7), and *Ae. searsii* (6), and *Ae bicornis* (6)) (Avni et al. 2022; Yang et al. 2023), which are the closest extant species to the donor of the B genome in the polyploid wheat species.

### Expression analysis of HIFs

Since one of the polymorphisms detected in the haplotype analyses was a 220-bp indel in the promoter region of *LFY-B*, we decided to study its effect on gene expression. We first selected plants homozygous for the indel alternative alleles from HIF40 derived from the Berkut x PB343 population. Grains were stratified for two days at 4 °C in the dark and then planted in growth chambers (PGR15, Conviron, Manitoba, Canada). Lights were set to 350 μmol m^-2^ s^-1^ at the canopy level. Plants were grown under inductive 16-h long days with temperatures set during the day to 22°C and during the night to 17°C. Shoot apical meristems were collected at the lemma primordia developmental stage (W3.25 in Waddington scale) (Waddington et al. 1983) and six apices were pooled for each of the six replicates collected from each genotype. RNA was extracted using the Spectrum Plant Total RNA Kit (Sigma-Aldrich, St. Louis, MO, USA). One μg of RNA was treated with RQ1 RNase-Free DNase (Promega, M6101) first and then used for cDNA synthesis with the High-Capacity cDNA Reverse Transcription Kit (Applied Biosystems, Foster City, CA, USA). The qRT-PCR experiments were performed using Quantinova SYBR Green PCR kit (Qiagen, 208052) in a 7500 Fast Real-Time PCR system (Applied Biosystems) with *LFY-B* genome specific primers (Table S1). For PCR reactions, we used one cycle at 95 °C for 2 min and 40 cycles of 95 °C for 5 s and 60 °C for 30 s, followed by a melting curve program. Transcript levels for all genes are expressed as linearized fold-ACTIN levels calculated by the formula 2^(ACTIN CT – TARGET CT)^.

### Epistatic interactions between natural variants of *LFY-B* and *WAPO-A1*

To evaluate the combined effect of the natural variants in *LFY-B* and *WAPO-A1*, we characterized two hexaploid and two tetraploid populations segregating for both genes. For *WAPO-A1*, we focused on the C47F amino acid change which was previously shown to be associated with significant differences in SNS (Kuzay et al. 2022). The first hexaploid population, Berkut x RAC875, included 75 recombinant inbred lines (RILs) that were previously evaluated for SNS in six different environments (Zhang et al. 2018). The second hexaploid population, RSI5 x RAC875, was evaluated in three separate experiments, first as F_2_, then as F_2:2_ plants derived from a single F_2_ plant heterozygous for *LFY-B* and *WAPO-A1*, and finally as F_3:2_ plants derived from a single F_3_ plant heterozygous for both genes. Plants homozygous for the four possible *LFY-B/WAPO-A1* allele combinations were selected with molecular markers and evaluated for SNS. Average SNS was calculated from the two tallest spikes in each plant.

For the first tetraploid population, we crossed Kronos x Gredho (which carried the desired *LFY-B* allele) and then backcrossed it to a Kronos line in which we had previously introgressed the *WAPO-A1* 47F allele from hexaploid wheat (Kuzay et al. 2022). We backcrossed this plant four times to the Kronos-47F line, selecting in each generation plants heterozygous for both genes using molecular markers. Finally, we generated BC_4_F_2_ seeds from a single BC_4_F_1_ plant heterozygous for both genes and sowed the grains in the UCD experimental field in November 2022 as single BC_4_F_2_ plants. The second tetraploid population was generated from the cross between Kronos and an unusual durum line from Syria (PI 519639) which carried the desired *LFY-B / WAPO-A1* allele combination. The F_1_ was backcrossed once to Kronos, and one BC_1_F_3_ plant heterozygous for both alleles was used to generate BC_1_F_3:2_ seeds that were sown in the same UC Davis experimental field side by side with the last population. We genotyped both populations using molecular markers for *LFY-B* and *WAPO-A1*, and evaluated the mature plants homozygous for the four different *LFY-B/WAPO-A1* combinations in 2023 for SNS (using three spikes per plant).

## Results

### Natural variants in *LFY* affect SNS in wheat

To test if *LFY* was associated with natural variation in SNS in polyploid wheat, we explored a set of eight wheat Nested Association Mapping (NAM) populations previously characterized for this trait (Zhang et al. 2018) and genotyped using the Infinium wheat SNP 90K iSelect assay (Illumina Inc., San Diego, CA, USA) (Wang et al. 2014). We did not find significant associations between SNS and SNP markers selected within 20-Mb regions encompassing *LFY-A* or *LFY-D*, but we did find significant associations between SNS and SNP markers flanking *LFY-B,* including IWB35244 as the peak marker (located 0.355 Mb proximal to *LFY-B* on chromosome arm 2BL) in the Berkut x PBW343 RIL population. Lines carrying the Berkut allele showed significantly higher SNS than those with the PBW343 allele across three field trials (Table S2). To validate this result, we generated heterogeneous inbred families (HIFs) from two F_5_ plants heterozygous for the *LFY-B* region. Sister F_5:2_ lines homozygous for the Berkut allele showed 1.6 to 1.8 more spikelets per spike than those homozygous for the PBW343 allele (*P* < 0.001, Fig. 1A, Table S3), confirming the significant effect of this chromosome region on spike development.

To test if the effect of this chromosome region on SNS was linked to *LFY-B* or not, we generated a high-density genetic map of a 3 Mb region on chromosome arm 2BL encompassing the *LFY-B* gene. We first developed molecular markers 2B-657.1 and 2B-660.1 flanking this region, and used them to screen 402 F_5:3_ plants derived from the Berkut x PBW343 HIFs (804 segregating chromosomes). This screening yielded 8 recombination events indicating a genetic distance of 1.0 cM, and a ratio between physical and genetic distances in this region of 3.1 Mb per cM.

We then developed molecular markers to characterize the lines with new recombination events within the candidate gene region. This task was facilitated by available exome capture data for the two parental lines in the T3/Wheat database (https://wheat.triticeaetoolbox.org/ dataset Exome_Capture_2017_WheatCAP_UCD). We first developed a KASP marker for a polymorphism located within the first exon of *LFY-B* (C-658199056-A) that resulted in a change from arginine to serine at position 80 (henceforth, R80S, Fig. 1B). We used these markers to genotype the recombinant lines, and selected three lines with the closest recombination events to *LFY-B* for phenotypic characterization. The field experiment showed significant differences in SNS between sister lines, which indicates that the causal gene of the SNS-QTL is located within the 760 kb region flanked by markers 1142/1143 (2B: 657,843,693) and 1148/1149 (2B: 658,603,716, Fig. 1C).

The candidate region for the SNS QTL includes only two complete and two partial high-confidence annotated genes in addition to *LFY-B* (Fig. 1D, Table S4). Among these genes, *TraesCS2B02G464000* and *TraesCS2B02G464100* encode proteins with kinase domains that are expressed at very low levels during spike development (VanGessel et al. 2022). *TraesCS2B02G464300* is highly expressed in the developing spike (VanGessel et al. 2022), but encodes a 50S ribosomal protein L11 that has no amino acid polymorphisms between the parental lines. Finally, only the promoter region of *TraesCS2B02G464400* is located within the candidate gene region, and it does not show major polymorphisms. By contrast, *LFY-B* is highly expressed in the developing spike (VanGessel et al. 2022), and shows a polymorphism in the conserved SAM domain (Fig. 2A) that can contribute to the observed differences in SNS. These results suggest that, among the few genes identified within the candidate region, *LFY-B* is the strongest candidate for the SNS QTL detected on chromosome arm 2B in the Berkut x PBW343 population.

**Fig. 2.**
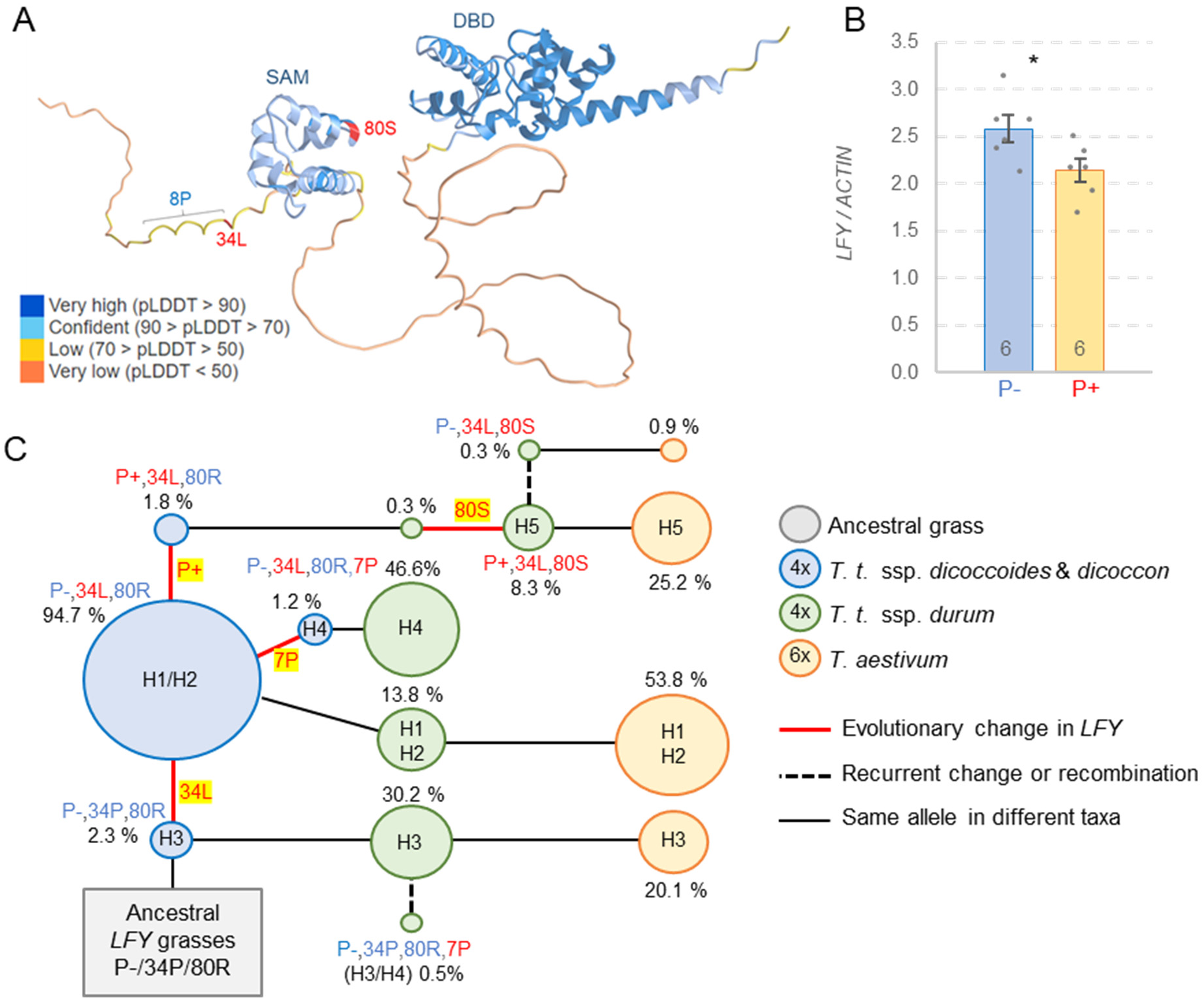
Natural variation in *LFY-B* in wheat. (**A**) Location of the natural polymorphisms (in red) in an AlphaFold simulation of the wheat LFY protein for haplotype H5. Colors indicate confidence levels (pLDDT = Local Distance Difference Test), SAM = Sterile Alpha Motif oligomerization domain and DBD = DNA binding domain. (**B**) Quantitative Reverse Transcription PCR (qRT-PCR) comparison of *LFY* expression between HIF sister lines homozygous for the H5 (P+ 80S) or H2 haplotypes (P-80R) in developing spikes at the lemma primordia stage. The number of biological replications is indicated at the base of the bars. Error bars indicate SEM and probability values are from two tailed *t-*tests. * = *P <* 0.05. Raw data and statistics are available in Table S7. (**C**) Changes in *LFY-B* allele frequencies in ancestral tetraploid (blue circles), modern tetraploid (green circles) and modern hexaploid (orange circles) wheat accessions. The likely origins of the different polymorphisms are marked in red bold lines and highlighted.

### *LFY-B* alleles are defined by multiple changes in promoter and coding regions

To better characterize the natural variation in the *LFY-B* region, we performed a haplotype analysis for an ∼800 kb region of chromosome arm 2BL including *LFY-B* (see Materials and Methods). The 146 SNPs identified in this region defined five haplotypes designated here as H1 to H5 (Table S5). We then determined the haplotypes present in this region in 17 sequenced wheat genomes (Avni et al. 2017; Maccaferri et al. 2019; Walkowiak et al. 2020; Sato et al. 2021), and used them to obtain the *LFY-B* sequences associated with each haplotype (Table S6). The H3 haplotype was not present among these sequenced wheat genomes, so we Sanger-sequenced *LFY-B* from hexaploid wheat cultivar RAC875 and deposited the sequence in GenBank (OR753911).

Comparison of the *LFY-B* coding sequences associated with the five different haplotypes revealed the previously described R80S polymorphism, a proline to leucine change at position 34 (P34L), and a proline deletion in a stretch of eight prolines at positions 26 to 33 (henceforth 8P to 7P). To better understand the potential effect of these amino acid changes, we looked at their position within the protein in an AlphaFold reconstruction of the LFY-B protein (Fig. 2A). The R80S polymorphism is located within the N-terminal conserved Sterile Alpha Motif oligomerization domain (SAM, Fig. 2A) (Siriwardana and Lamb 2012; Sayou et al. 2016), and has a negative BLOSUM 62 score of -1 suggesting potential effects on the structure and/or function of the LFY-B protein. Although the 8P to 7P and P34L polymorphisms are located outside the conserved SAM and DNA binding domain (DBD) region, they are still located within a region of LFY that is well conserved among grass species (Fig. S1).

A comparison of the *LFY-B* promoter region revealed the presence of a 220-bp insertion located 872 to 653 bp upstream from the *LFY-B* start codon that was associated with the H5 haplotype. The presence of this insertion is indicated hereafter as P+ and its absence as P- (Fig. 1B). To estimate the potential effect of the promoter indel, we compared *LFY-B* expression levels in HIFs carrying the Berkut (H5, P+) and PBW343 (H2, P-) haplotypes. The H5 haplotype was associated with a 17 % decrease in *LFY-B* transcript levels in the developing spikes at the lemma primordia stage relative to H2 (*P=* 0.036, Fig. 2B, Table S7). However, since the 220-bp insertion is not the only polymorphism that differentiates the H5 and H2 haplotypes, this data is insufficient to demonstrate that the differences in *LFY-B1* expression in the developing spike between the H2 and H5 haplotypes are caused by the promoter indel.

### *LFY-B* haplotype frequencies changed during wheat domestication

The LFY-B allele combining the 34P, 80R, and 8P sequences (H3 haplotype, Fig. 1B) is likely the ancestral haplotype because it is present in most other grass species (Fig. S1). The derived haplotypes H1 and H2 haplotypes differ from H3 by the presence of the 34L polymorphism (P-,34L,80R,8P). This polymorphism is also present in the *LFY-B* alleles associated with the H4 and H5 haplotypes, but in the H4 haplotype it is combined with the proline deletion (P-,34L,80R,7P) and in the H5 haplotype with the 80S polymorphism and the promoter insertion (P+,34L,80S,8P) (Fig. 1B).

To estimate the frequency of the different *LFY-B* changes, we designed markers for the promoter insertion, the two non-synonymous amino acid changes (P34L and R80S) and the deletion of the proline in the 8P to 7P polymorphism (primers in Table S1), and used them to characterized 171 wild and 618 cultivated wheat accessions (Table S8). We found the ancestral *LFY-B* allele P-,34P,80R (H3 haplotype) in 30.2% of the *T. turgidum* ssp. *durum* and 20.1 % of the *T. aestivum* accessions; but surprisingly in only 2.3 % of the ancestral tetraploid species *T. turgidum* ssp. *dicoccoides* and *T. turgidum* ssp. *dicoccon* (Fig. 2C and Table S8). The *LFY-B* allele P-,34L,80R found in the H1 and H2 haplotypes was the most abundant in all wheat taxa (Fig. 2C), suggesting that the 34L change was an early event in wheat evolution that was favored by selection. We did not find the R80S polymorphism among the ancestral tetraploid accessions, but found it in 8.3% of the modern durum and in 25.2 % of the common wheat varieties, suggesting positive selection. Among the accessions with the 80S polymorphism, 96.8% also carry the 220-bp promoter insertion. Finally, we detected the 8P to 7P deletion associated with the H4 haplotype in few ancestral tetraploids (1.2 %), at a higher frequency in modern durum (46.6 %) accessions, and absent in hexaploid wheat (Fig. 2C).

### Natural alleles of *LFY-B* and *WAPO-A1* show significant genetic interactions for SNS

Loss-of-function alleles *lfy* and *wapo1* show reciprocal recessive epistatic interactions for SNS, indicating that functional copies of both genes are required to increase SNS. Therefore, we explored if the natural variants of *LFY-B* discovered in this study were able to interact with the *WAPO-A1* alleles reported in previous studies (Kuzay et al. 2022; Kuzay et al. 2019) in the regulation of SNS. We evaluated these interactions in four different populations segregating for the *WAPO-A1* C47F polymorphism and for different *LFY-B* alleles.

In the Berkut (H5, 47F) x RAC875 (H3, 47C) population, the parental *LFY-B* alleles shared the 8P sequence but differed in the 220-bp promoter indel and the P34L and R80S polymorphisms. A combined ANOVA for the six different year/location combinations where this population was evaluated (Zhang et al. 2018) showed highly significant effects on SNS for both *WAPO-A1* (*P* < 0.001) and *LFY-B* (*P* = 0.0028), as well as a significant interaction between the two genes (*P* = 0.0013) (Table S9). The effect of *LFY-B* on SNS was significant only in the presence of the 47F allele, where plants carrying the H5 haplotype had 1.4 more spikelets per spike than plants with the H3 haplotype (*P* < 0.001, Fig. 3A). The effect of *WAPO-A1* was significant in the presence of both *LFY-B* alleles, but was stronger in the presence of the H5 haplotype (2.5 spikelets/spike) than in the presence of ancestral the H3 haplotype (1.0 spikelets per spike, Fig. 3A, Table S9).

**Fig. 3.**
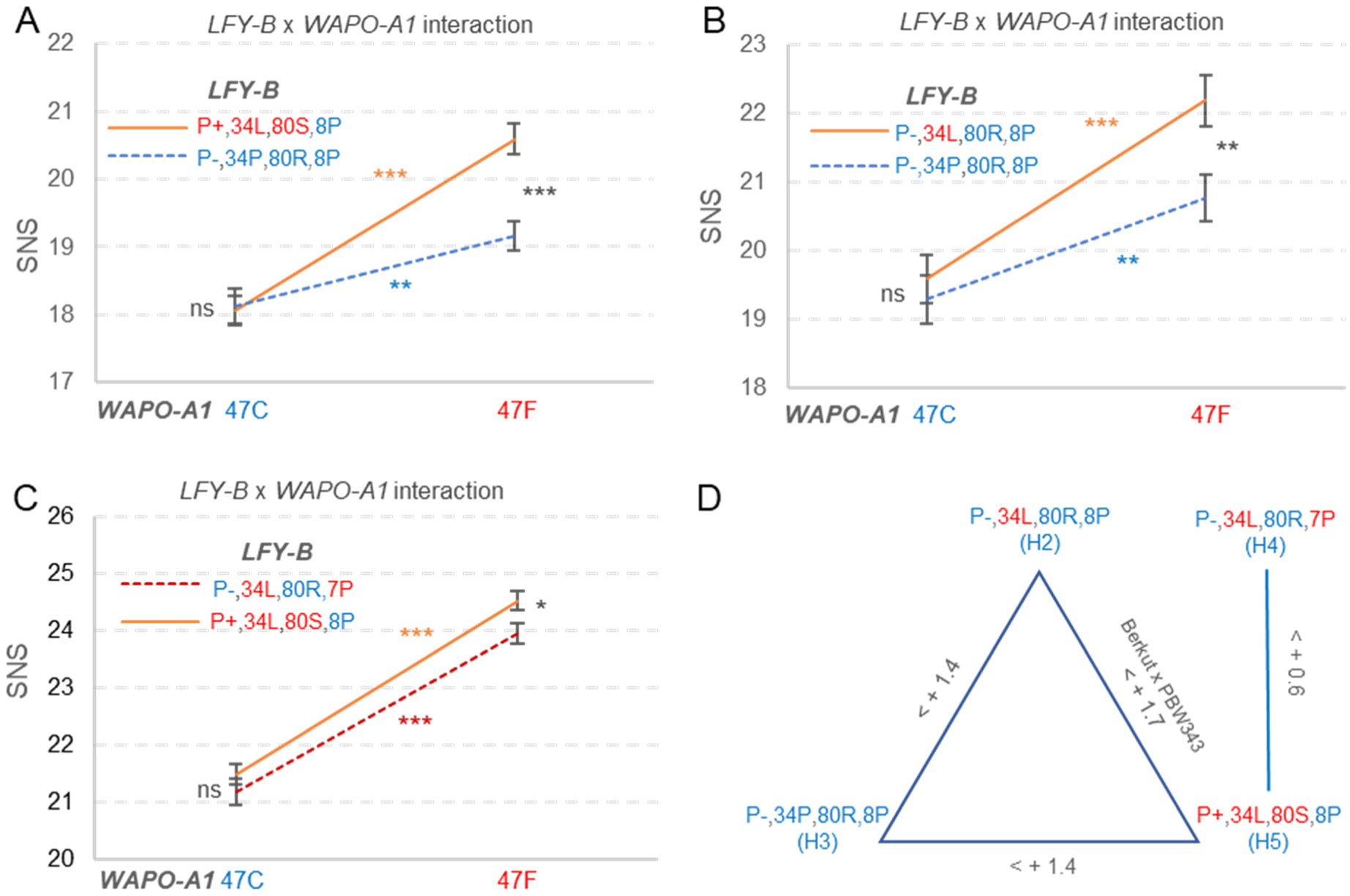
Genetic interactions for SNS between *LFY-B* and *WAPO-A1* natural alleles. Interactions between the *WAPO-A1* C47F alleles and different *LFY-B* alleles. (**A**) Berkut (H5) x RAC875 (H3) RIL population evaluated in 6 environments, total N = 390 (Zhang et al. 2018). (**B**) RSI5 (H2) x RAC875 (H3). The statistics are from the combined ANOVA of three experiments using experiment as block, total N = 118. (**C**) Combined analysis of tetraploid populations Kronos (H4) x PI 519639 (H5) BC_1_F_3:2_and Kronos-47F (H4) x Gredho (H5) BC_4_F_2_ evaluated in the field. *P* values indicate significance of the four possible contrasts in the ANOVAs described in Table S9. (**D**) Summary of the effects of different *LFY-B* alleles in the *WAPO-A1* 47F background. ns = not significant, * = *P <* 0.05, ** = *P <* 0.01, and *** = *P <* 0.001. Error bars are SEM. Statistics are available in Table S9.

In the second population, RSI5 (H2, 47F) x RAC875 (H3, 47C), the parental *LFY-B* alleles shared the P-/80R/8P sequences, and differed in the P34L alleles. This population showed significant effects on SNS for both *WAPO-A1* (*P* < 0.001) and *LFY-B* (*P* = 0.0165). The interaction was not statistically significant (*P* = 0.1142) but, as in the previous population, *LFY-B* showed a stronger effect on SNS in the presence of the *WAPO-A1* 47F allele (1.4 spikelets/spike, *P =* 0.0056) than in the presence of the 47C allele (0.3 spikelets/spike, *P =* 0.5481, Fig. 3B, Table S9).

In the last two populations, Kronos (H4, 47C) was crossed with two different tetraploid lines carrying the H5, 47F allele combinations (see Materials and Methods). These two *LFY-B* haplotypes share the 34L polymorphisms but differ in the promoter insertion and the R80S and 8P to 7B polymorphisms. Since the two populations segregated for the same alleles, we combined the statistical analysis using population as a block (Table S9). The P+,80S,8P allele was associated with a significant increase in SNS (*P* = 0.0205), which was stronger in the presence of the *WAPO-A1* allele 47F (0.6 spikelets/spike, *P* = 0.0190) than in the presence of 47C, where no significant differences were detected (*P =* 0.2892, Fig. 7D). The differences between *WAPO-A1* 47F and 47C alleles were significant in the presence of both *LFY-B* alleles (*P* < 0.0001, Fig. 7D).

Figure 3D summarizes the effects of the different *LFY-B* alleles in the presence of the *WAPO-A1* 47F allele from Figs. 3A-C, and from the original HIFs segregating for the Berkut (H5, 47F) and PBW343 (H2, 47F) alleles (Fig. 1A and combined analysis in Table S9). In summary, in all three populations the positive effects of the derived *LFY-B* alleles on SNS relative to the more ancestral alleles were significant only in the presence of the *WAPO-A1* 47F allele (high SNS).

The largest beneficial effect on SNS was observed for the H5 haplotype, which combines the 220-bp promoter insertion with the 34L and 80S polymorphisms.

## Discussion

### Natural variation in *LFY-B* is responsible for the SNS QTL detected in this region

The high-density map of the SNS QTL on chromosome 2BL narrowed down the candidate gene region to 770 kb, a region including only two complete and two partial genes in addition to *LFY-B.* These additional genes were expressed at very low levels in the developing spikes (VanGessel et al. 2022) or showed no clear polymorphisms between the two parental lines. By contrast, *LFY-B* is highly expressed in the developing spikes (Paraiso et al. 2024) and shows two polymorphisms between the parental lines that affect its expression levels (Fig. 2B) and the encoded protein. The R80S amino acid polymorphism is located within the conserved SAM domain that plays an important role in LFY oligomerization (Sayou et al. 2016) (Fig. 2A). In Arabidopsis, a T75E mutation in the SAM domain disrupts LFY oligomerization and affects access to low affinity sites or closed chromatin. (Sayou et al. 2016). That conserved threonine is located at position 73 in *LFY-B* in wheat, only seven amino acids from the R80S polymorphism. In addition to its location in a conserved domain, the wheat R80S mutation has a negative BLOSUM 62 score (-1) that is indicative of a potential disruptive effect on protein structure or function.

The previous results suggest that *LFY-B* is the most promising candidate for the SNS QTL among the five genes detected in the candidate gene region. This hypothesis is also supported by the 8% reduction in SNS (*P* = 0.003) observed in an EMS-induced truncated mutant of *LFY-B* (Q249*) in tetraploid wheat Kronos (Paraiso et al. 2024), and the even stronger reduction observed in the combined *lfy-A lfy-B* mutant. This last result indicates that both homoeologs positively and redundantly regulate SNS in wheat and that a functional *LFY* gene is required to generate normal SNS. Changes in *LFY* expression are also sufficient to affect SNS. Overexpression of *LFY* under the maize ubiquitin promoter increased SNS, and partially complemented the effects of the loss-of-function mutations (Paraiso et al. 2024). Taken together, the high-density mapping results presented in this study, and the *LFY* mutants and transgenic results presented by Paraiso et al. (2024) indicate that *LFY-B* is the causal gene for the SNS QTL detected in the Berkut x PBW343 population.

### Rapid frequency increases of the derived *LFY-B* alleles suggest positive selection

The role of *LFY* on the regulation of SNS is also reflected in the association of different *LFY-B* natural alleles with differences in SNS. The P34L polymorphism was associated with significant differences in SNS in the RSI5 (H2) x RAC875 (H3) population (Fig. 3B). The *LFY-B* alleles in these two haplotypes share the P-,80R,8P sequences, so the 7% increase in SNS (in the 47F *WAPO-A1* background) is likely associated with the presence of the 34L polymorphism. The beneficial effect of the 34L polymorphism is also supported by its high frequency among the wild and cultivated emmer accession (Fig. 2C, 94.7%).

In spite of its high frequency in the polyploid wheat species, 34L is likely the derived allele because the 34P allele is highly conserved among other grass species (Fig. S1). We also found the 34P allele in 35 accessions of *Ae. speltoides* (9), *Ae. sharonensis* (7), *Ae. longissima* (7), and *Ae. searsii* (6), and *Ae bicornis* (6) (Table S8) (Avni et al. 2022; Yang et al. 2023), which are part of the Section Sitopsis where the B genome was originated. This result shows that the 34P allele was predominant in the ancestors of the B genome, and suggests that the 34L allele possibly originated in the wild tetraploid species. The high frequency (Fig. 2C) and wide geographic distribution (Table S8) of the 34L allele among the wild and cultivated emmer suggests that this allele originated early in the history of tetraploid wheat and was favored by positive selection. The higher frequency of the 34L among the modern hexaploid (80%) than among the durum varieties (70%), may have been favored by the higher frequency of the 47F allele among the hexaploid (>60%) than among the durum and cultivated emmer (<5%) (Kuzay et al. 2019) and the stronger effect of the 34L polymorphism in the presence of the *WAPO-A1* 47F allele compared to the 47C allele.

The P+,80S polymorphism (H5) showed positive and significant effects on SNS relative to the H2 (Fig. 1A), H3 (Fig. 3A) and H4 (Fig. 3C) haplotypes, which were larger in the presence of the *WAPO-A1* 47F allele. However, there might be additional epistatic interactions modulating the effect of the P+,80S allele, since we failed to detect significant differences in SNS in two of the six NAM populations segregating for this allele in spite of the presence of the 47F *WAPO-A1* allele (Zhang et al. 2018). The beneficial effect of the P+,80S is also supported by its rapid increase in allele frequencies. Since this polymorphism was not detected in the wild or cultivated emmer, we speculate that it originated more recently in cultivated durum or common wheats, which were domesticated ∼10,000 years ago (Dubcovsky et al. 2007). The faster increase in the H5 frequency among modern hexaploid cultivars (25.2%) relative to tetraploid wheat (8.3%, Fig. 2C) may be explained by the stronger effect of H5 on SNS in the presence of the 47F than the 47C *WAPO-A1* alleles (Fig. 3A), and the higher frequency of the *WAPO-A1* 47F allele in hexaploid wheat mentioned above.

Among the 98 accessions carrying the 80S or P+ polymorphisms (Table S8), 93% carry both alleles, suggesting that one originated within the other one soon after its origin, or that this combination has been favored by positive selection. We found the two alternative combinations, P+,80R (three *T. turgidum* ssp. *dicoccoides* and one durum) and P-,80S (one durum and two hexaploid), suggesting that one of these lineages is likely the result of recombination, gene conversion or recurrent mutations. We favor the hypothesis that the P+ insertion originated first in wild wheat and that the 80S polymorphism originated in a durum or common accession carrying the P+ insertion (Fig. 2C), but we cannot rule out alternative scenarios. These low-frequency alternative combinations provide useful tools to test if the beneficial effect of the H5 haplotype on SNS is associated with the 80S polymorphism, the 220-bp promoter insertion or both.

None of the populations presented in this study tested the effect of the 8P to 7P polymorphism on SNS separately from other *LFY-B* polymorphisms. However, a comparison of the differences in SNS between the H5 and H2 haplotypes (1.7 spikelets, Fig. 1A) and between the H5 and H4 haplotypes (0.6 spikelets Fig. 3C-D) provides an indirect estimate of the 8P to 7P effect. We speculate that the smaller effect of the P+,80S on SNS relative to H4 than to H2 may be caused by a small beneficial effect of the 7P allele on SNS, which would reduce the difference between H5 and H4. If this hypothesis is correct, it may explain the rapid increase of the frequency of the 7P polymorphism among the commercial tetraploid wheat varieties (Fig. 2C). However, the previous comparisons involve experiments performed under different conditions and, therefore, will need to be validated in more controlled experiments.

### Practical applications of the epistatic interactions among *WAPO-A1* and *LFY-B* natural alleles for SNS

The increases in the frequencies of the alleles associated with higher SNS in *LFY-B* described above parallel those in *WAPO-A1* described previously (Kuzay et al. 2019), and possibly reflect the synergistic effects among these alleles on SNS. For example, we only detected a significant effect for the P+,80S polymorphisms when the *WAPO-A1* 47F allele was present (Fig. 3A).

Similarly, the positive effect of the 47F allele on SNS was stronger in the presence of the *LFY-B* 34L or P+,80S alleles (Fig. 3A-B). The synergistic interaction of the *LFY-B* and *WAPO-A1* natural alleles in the regulation of SNS has likely favored the simultaneous selection for the most favorable combinations. The relatively fast increase in the frequencies of the beneficial *LFY-B* and *WAPO-A1* allele combinations in modern wheats suggests that breeders’ selection for grain yield or for larger spikes has been effective to drive these changes.

The identification of optimal *LFY-B - WAPO-A1* allele combination provides wheat breeders new tools to improve SNS. Markers for the favorable alleles can be used to implement marker assisted selection (MAS) programs, and the SNPs associated with the beneficial alleles can be incorporated as fixed factors into the genomic selection models to improve their efficiency (Sarinelli et al. 2019; Spindel et al. 2016). Among the common wheat accessions that already have the 47F allele (>60 %, (Kuzay et al. 2019), it is sufficient to incorporate the beneficial *LFY* alleles that work well with 47F (34L and P+,80S). However, in common wheat varieties lacking the 47F allele, the best strategy would be to simultaneously introgress the optimal *LFY-B - WAPO-A1* allele combinations. The incorporation of these beneficial combinations represents a particularly interesting opportunity for tetraploid wheat, where both *WAPO-A1* 47F and *LFY-B* P+/80S alleles are rare (Fig. 2C). It is tempting to speculate that the need to introgress both beneficial alleles simultaneously has acted as a barrier for the selection of these beneficial alleles in durum wheat. However, it is also possible that other epistatic interactions with the durum and common wheat genomes limit the effectiveness of these alleles in tetraploid wheat.

Pleiotropic effects are likely frequent among genes that encode proteins that physically interact with each other, since the function of the protein complex is likely affected by mutations in both partners. Another example of physically interacting wheat proteins showing genetic interactions for SNS was described recently for the natural alleles of FT-A2 and bZIPC1 (Glenn et al. 2023; Glenn et al. 2022). The alleles associated with improved SNS (*FT-A2* A10 and *bZIPC-B1* 151K-166M) showed increased frequencies during wheat domestication, and their combination resulted in synergistically positive effects on SNS (Glenn et al. 2023).

The gene regulatory networks that control spike development and grain yield components are highly interconnected to facilitate the incorporation of multiple environmental signals while maintaining balanced developmental processes. Therefore, frequent epistatic interactions are expected among the genes controlling these traits. A better understanding of these epistatic interactions would allow the identification of better allele combinations, and pave the way for future efforts to engineer more productive wheat varieties.

## Statements & Declarations

## Acknowledgments

We would like to thank Dr. Josh Hagerty for his help to manage the field experiments.

## Author contributions

JZ did the majority of the experimental work and data analysis and wrote the first draft. GFB ran the AlphaFold prediction for LFY protein. FP contributed to the expression experiments. JZ and JD conceived the project. JD obtained funding and administered the project, contributed data analyses, supervised JZ, FP and GFB, and was responsible for the final manuscript. All authors reviewed the manuscript and provided suggestions.

## Data availability

All data are available in the main text or the supplementary materials. The sequence of LFY for the H3 haplotype has been deposited in GenBank as accession OR753911.

## Ethics approval

Do not apply to this study.

