## Supplementary Figure 1 for "Natural alleles of *LEAFY* and *WAPO1* interact to regulate spikelet number per spike in wheat"

**Figure S1. Alignment of LFY proteins from multiple grass species**. The query is the *T. aestivum LFY-B* haplotype 5 (P+/34L/80S). Only the region around the L34P and R80S polymorphisms is presented. *Triticum urartu* is included as the *T. aestivum* donor of the A genome and *Aegilops tauschii* as the donor of the D genome. These two species as well as several other grasses carry the 34P and 80R alleles, which are considered as the ancestral alleles. Underlined is a stretch of eight proline residues that are reduced to seven in haplotype H4 (8 P to 7 P).

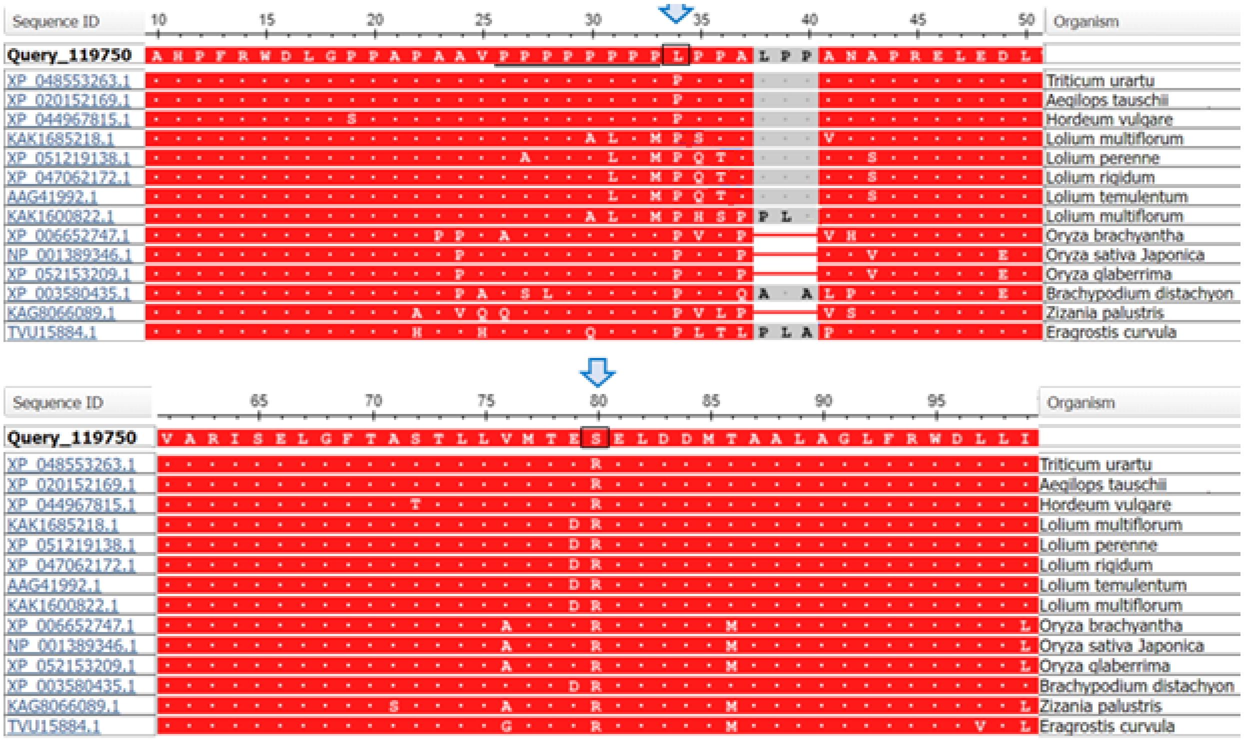
